# Quantitative Phenotyping of Vascular Damage Caused by Fusarium Wilt Disease in Cowpea

**DOI:** 10.1101/850701

**Authors:** Arsenio D. Ndeve, Philip A. Roberts

**Affiliations:** University of California, Riverside, CA 92521, USA

**Keywords:** *Fusarium oxysporum* f. sp. *tracheiphilium*, vascular wilt, *Vigna unguiculata*, race-specific resistance

## Abstract

Assessment of the severity of Fusarium wilt disease in cowpea and other crops relies mainly on visual rating scales which are prone to errors, which can compromise the reproducibility of the data. Furthermore, the rating scales require considerable practical training and routine experience for reliable assessment. Two objective metrics, stem vascular discoloration length (%VDL) and number of Fusarium necrotic vessels (NFNV), for quantitative measurement of vascular damage incited by *Fusarium oxysporum* f. sp. *tracheiphilum* race 4 (Fot4) of cowpea, were compared and their utility as a measure of disease severity and potential usefulness in other crop pathosystems is proposed. The metrics were tested in seven F_2_ populations and one F_2:3_ population, segregating for wilt response, and inoculated with race Fot4 at the seedling stage. %VDL and NFNV were highly correlated with plant wilting for all populations (*r =* 0.51 – 0.93 and 0.52 – 0.94, respectively). Furthermore, the relationships between the variables were linear in all populations (*R*^*2*^ = 0.81 to 0.87 and 0.71 to 0.91), indicating that they can provide accurate and reliable measurement of severity of Fusarium wilt disease. Also, %VDL and NFNV were strongly correlated (*r =* 0.88 - 0.97) and demonstrated a linear relationship (*R*^*2*^ = 0.69 – 0.94). Analysis of goodness-of-fit in two F_2_ populations revealed that errors in measurement of vascular discoloration length can result in higher segregation distortion when compared to enumeration of necrotic vessels. However, both metrics were highly effective in accounting for the severity of vascular damage caused by Fusarium wilt disease.

---

*Fusarium oxysporum* f. sp. *tracheiphilum* (E. F. Sm) Snyder and Hansen is the causal agent of wilt disease in cowpea, resulting in poor stand establishment, poor plant growth (Smith et al., 1999), and resulting yield loss up to 65% (Singh, 2014). The destructive effect of Fusarium wilt on cowpea production justifies research to understand and ameliorate its impact, and for breeding for resistance to the disease. The identification of an effective source of Fusarium wilt resistance, the development of segregating breeding populations and selection of superior breeding lines carrying resistance alleles, coupled with genomic analysis are crucial steps for breeding Fusarium wilt resistant cultivars. Phenotype-based screening for resistance donors and selection of superior breeding lines requires a sensitive screening method with reliable metrics to accurately assess the severity of disease. Reliable disease-phenotyping metrics allow for accurate identification of true phenotypic disease responses (resistant, moderately resistant and susceptible) in screening assays (Luckew et al., 2012). Fusarium wilt resistant and susceptible cowpea genotypes can be easily identified in a germplasm collection or in a segregating population when the trait is under control by one or two genetic factors. However, under quantitative inheritance effective selection for resistant genotypes can be compromised by the lack of an objective screening method; therefore, in these cases quantitative metrics can provide accurate and precise quantification of the severity of disease (Ndeve, 2017). In resistant and moderately resistant genotypes, disease severity expressed as the extent of plant wilting can be weakly associated with length of vascular discoloration which is an indicator of the extent of vascular colonization and the damage caused by the fungus (Talboys, 1972; Luckew et al., 2012; Ndeve, 2017). Less objective disease assessment is error prone and can lead to inaccurate association between plant wilting and the extent of vascular discoloration and damage, which could compromise the individual usefulness of each metric in Fusarium wilt disease severity assessment and the translation of the results into practical use.

Variation in wilt disease phenotypes has been reported in several studies. In common bean and cotton, extensive and severe colonization of the vascular system occurs on susceptible cultivars and breeding lines, and plant wilting is strongly correlated with vascular discoloration (Pastor Corrales and Abawi, 1987; Buruchara and Camacho, 2000; Ulloa et al., 2006). In both crops, some resistant common bean cultivars and cotton breeding lines exhibited mild wilting and discoloration of vascular tissues. In a study by Gao et al. (1994) on eight tomato cultivars screened for resistance to five forms and races of *Fusarium oxysporum* f. sp. *lycopersici*, vascular discoloration was strongly correlated with plant wilting, and there was no evidence of a tolerant reaction. According to Talboys (1972), Fusarium wilt tolerant cultivars can be misclassified as resistant cultivars if disease severity is quantified solely based on the external phenotypic wilt reaction because systemic vascular colonization in tolerant cultivars does not necessarily lead to plant wilting. As in cowpea, susceptible common bean and cotton cultivars can display wilt symptoms 5 to 9 days after inoculation, and severe or complete plant wilting and defoliation, which indicates the severity of damage to the vascular system, can occur 14 days after inoculation (Pastor Corrales and Abawi, 1987; Ulloa et al., 2006). Luckew et al. (2012) reported that in lines of soybean from resistant × resistant crosses, the severity of wilt caused by *Fusarium virguliforme* was not correlated with the severity of damage in the root system. These reports indicate that reliance on plant wilting to quantify severity of Fusarium wilt and the use of qualitative and semi-quantitative metrics can compromise the accuracy of wilt disease severity estimation, especially if plant wilt phenotypes are not highly associated with vascular discoloration symptoms as the two traits are under control by distinct mechanisms (Talboys,1972; Luckew et al., 2012)

Plant disease severity assessment in many crop species relies mostly on visual assessment based on rating scales (Bock and Nutter, 2011; Chiang et al., 2014). In general, severity of fusariosis is quantified utilizing rating scales in which the external plant phenotypic response (wilting) is used to express the severity of disease. Disease assessment rating scales include ordinal scales of 1 – 5 (Rigert and Foster, 1987) 1 – 9 (Van Schoonhoven and Pastor Corrales, 1987; Pastor Corrales and Abawi, 1987), nominal or descriptive scales (resistant and susceptible) (Lv et al., 2014) and nearest percent estimates or ratio scale (Walker, 1930; Palti and Joffe, 1971; Joffe et al., 1974; James, 1974; Netzer et al., 1977; Netzer and Weintall, 1980; Sharma et al., 2004; Lv et al., 2014; Chiang et al., 2014). Similarly to nominal scales, ordinal scales are also descriptive; however, the severity of disease is graded using arbitrary classes which indicate distinct levels of severity of disease symptoms (Bock et al., 2010).

Other Fusarium wilt disease severity assessment methods have employed estimates of both plant wilt and vascular discoloration phenotypes by using ordninal rating scales of 0 - 5 (Ulloa et al. 2006; Pottorff et al., 2012; Pottorff et al., 2014) or 1 – 9 (adapted from Pastor Corrales and Abawi, 1987) (Fall et al., 2001). However, Pastor Corrales and Abawi (1987) and Buruchara and Camacho (2000) used an ordinal rating scale of 1 – 9 to phenotype plant wilting and a nominal or descriptive scale to estimate extent of vascular discoloration. The severity of vascular damage was described as none, light, intermediate and severe. In cotton, Wang et al., 2017 assessed the severity of Fusarium wilt disease using an ordinal scale of 0 – 5 to express the severity of plant wilt and a ratio scale (0 – 100%) to express the extent of vascular discoloration.

Each of the visual rating scales described above is prone to errors which can compromise the accuracy, precision, repeatability and reproducibility of estimates of Fusarium wilt severity. Errors can limit correct differentiation among resistant, moderately resistant and tolerant plant phenotypes. Indeed, when less experienced individuals use visual rating scales to estimate disease severity, the magnitude of inaccuracy and imprecision in disease severity estimates can be substantial due to evaluator subjectivity, compromising the usefulness of the results for critical research or breeding decisions (Nutter et al., 1993; Bock et al., 2008).

This study introduces a novel, objective metric based on the number of necrotic vessels for phenotyping the extent of vascular symptoms incited by Fusarium wilt in cowpea. The relative accuracy of the new metric in expressing severity of wilt is contrasted with vascular discoloration length (%VDL), also an objective metric used to quantify the extent of vascular colonization by Fusarium wilt. The relationships between plant wilting symptoms and the objective metrics are explored. The concept accuracy is defined as the degree of closeness of an estimate to the true or actual value (Nutter et al., 1991), in this study for analytical purpose the true or actual phenotypic value is the plant wilt phenotype, and it is potentially predicted based on the severity of vascular discoloration or the number of necrotic vessels. Therefore, these two parameters are contrasted to determine to what extent their measured values closely reflect the severity of plant wilt (the true value).

## MATERIALS AND METHODS

### Plant material

One F_2:3_ and seven F_2_ cowpea populations were used to determine the variability in plant wilting (expressed as a wilt score), vascular discoloration length (expressed as the percent of stem height, %VDL), and vascular necrosis (expressed as number of Fusarium necrotic vessels - NFNV) phenotypes.

In brief, to generate the F_2_ and F_2:3_ populations, a Fusarium wilt race 4 (Fot4) resistant genotype (Parent A, Table 1) was crossed with three other resistant genotypes (Parents C, E and H; Table 1) to derive F_1_ generations of resistant × resistant populations (populations 5, 6 and 7), and with four susceptible genotypes (Parents B, D, F and G; Table 1) to derive F_1_ generations of susceptible × resistant populations (populations 1, 2, 3 and 4). In all crosses, parent A was used as the pollen donor male and all other parents as female. A single F_1_ seed of each cross was planted to generate seven F_2_ populations (Table 1); in addition, 175 seeds of F_2_ population 1 (Table 1) were planted to generate an F_2:3_ population (population 8, Table 1). The populations, their sizes and number of plants evaluated per parent (control) are presented (Table 1). The number of plants evaluated per F_2:3_ family varied from 8 to 31 (the average number of plants evaluated per family was 25).

**TABLE 1.** Cowpea population designations, crosses, population size, and number of plants per parent (control) evaluated in the Fusarium wilt assays.

| Population |  |  |  | Parents (controls) |  |  |  |  |  |  |  |
| --- | --- | --- | --- | --- | --- | --- | --- | --- | --- | --- | --- |
|  |  |  |  | Number of Plants |  |  |  |  |  |  |  |
| N<sup>o</sup> | Cross | Gen | Size | A | B | C | D | E | F | G | H |
| 1 | CB46/FN-2-9-04 | F<sub>2</sub> | 364 |  | 39 |  |  |  |  |  |  |
| 2 | INIA-73/FN-2-9-04 | F<sub>2</sub> | 353 |  |  |  | 28 |  |  |  |  |
| 3 | 24-125B-1/FN-2-9-04 | F<sub>2</sub> | 209 |  |  |  |  |  | 35 |  |  |
| 4 | Bambey-21/FN-2-9-04 | F<sub>2</sub> | 120 | 20 |  |  |  |  |  | 16 |  |
| 5 | CB27/FN-2-9-04 | F<sub>2</sub> | 323 |  |  | 18 |  |  |  |  |  |
| 6 | IT93K-503-1/FN-2-9-04 | F<sub>2</sub> | 363 |  |  |  |  | 21 |  |  |  |
| 7 | Ecute/FN-2-9-04 | F<sub>2</sub> | 145 |  |  |  |  |  |  |  | 25 |
| 8 | CB46/FN-2-9-04 | F<sub>2:3</sub> | 175 | 10 | 22 |  | 27 |  | 5 | 6 | 12 |

N<sup>o</sup> = population number; Gen = generation; parents A, B, C, D, E, F, G, H = FN-2-9-04, CB46, CB27, INIA-73, IT93K-503-1, 24-125B-1, Bambey-21 and Ecute, respectively.

The F_2_ populations and an F_2:3_ population were used for this study because of their high levels of segregation compared to late generation recombinant inbreeding lines. The levels of segregation in these populations were used to capture a full range of expected phenotypic responses (resistant, intermediate and susceptible) in cowpea screening assays; for example, in cowpea germplasm screenings aimed at identifying novel sources of resistance to Fot4, and in segregating populations carrying potential Fot4 resistance allele combinations. In cowpea breeding programs, phenotyping for Fot4 resistance in earlier generations (e.g. F_2:3_) is crucial for early plant selection and to capitalize on genotyping resources for effective and efficient plant selection.

### Inoculum preparation

A dried culture of Fot4 isolate T97-30 (isolated from infected cowpea plants in Bakersfield, California), derived from a single spore line was stored at −80 °C on potato dextrose agar (PDA) plates (Pottorff et al., 2014). The dried culture was prepared by culturing the fungus in a shallow and thin layer of PDA in a petri dish. A single 0.5-cm^2^ plug was cut from the dried culture and transferred onto a new petri dish containing fresh PDA amended with 3 mM streptomycin/liter. The culture was incubated at room temperature for 3-4 days under a 16 photoperiod. A fresh 1-cm^2^ plug was aseptically cut from the new culture and placed in a 500 ml Erlenmeyer flask containing freshly prepared and cooled potato-dextrose broth. The flask was incubated in a shaker for 4 days at 30 rpm, and 27 °C under constant light. The broth was filtered through 8 layers of cheesecloth, and the flow-through solution containing spores was collected in a beaker. The spores were counted using a hemocytometer under a light microscope, and the concentration was adjusted to 10^6^ microconidia/ml using sterile distilled water. The inoculum was used immediately for plant inoculation.

### Growth conditions and infection assays

The experiments were conducted in a controlled environment greenhouse at the University of California, Riverside (29 to 32 oC) under a 14 hours photoperiod. The F_2_, F_2:3_ and control genotypes were planted in seedling-trays containing Sungro^®^ Horticulture Sunshine mix #2 growing medium (Sun Gro Horticulture Canada Ltd). Following a modified protocol of Rigert and Foster (1987), seven days after planting, the seedlings were uprooted, and the root system washed, clipped to 3 cm length, dipped for 4 minutes into a fresh spore suspension of Fusarium wilt race four (Fot4) containing 10^6^ spores/ml water prepared as described above, and transplanted into 0.95 L foam cups (Fig. 1B and 1C) containing UC-Mix 3 soil (University of California Riverside, http://agops.ucr.edu/soil). The seedlings were watered once per day, and two weeks after inoculation they were fertilized using Osmocote Classic 14-14-14 fertilizer (Everris NA Inc., Dublin, OH).

**Fig.1.**
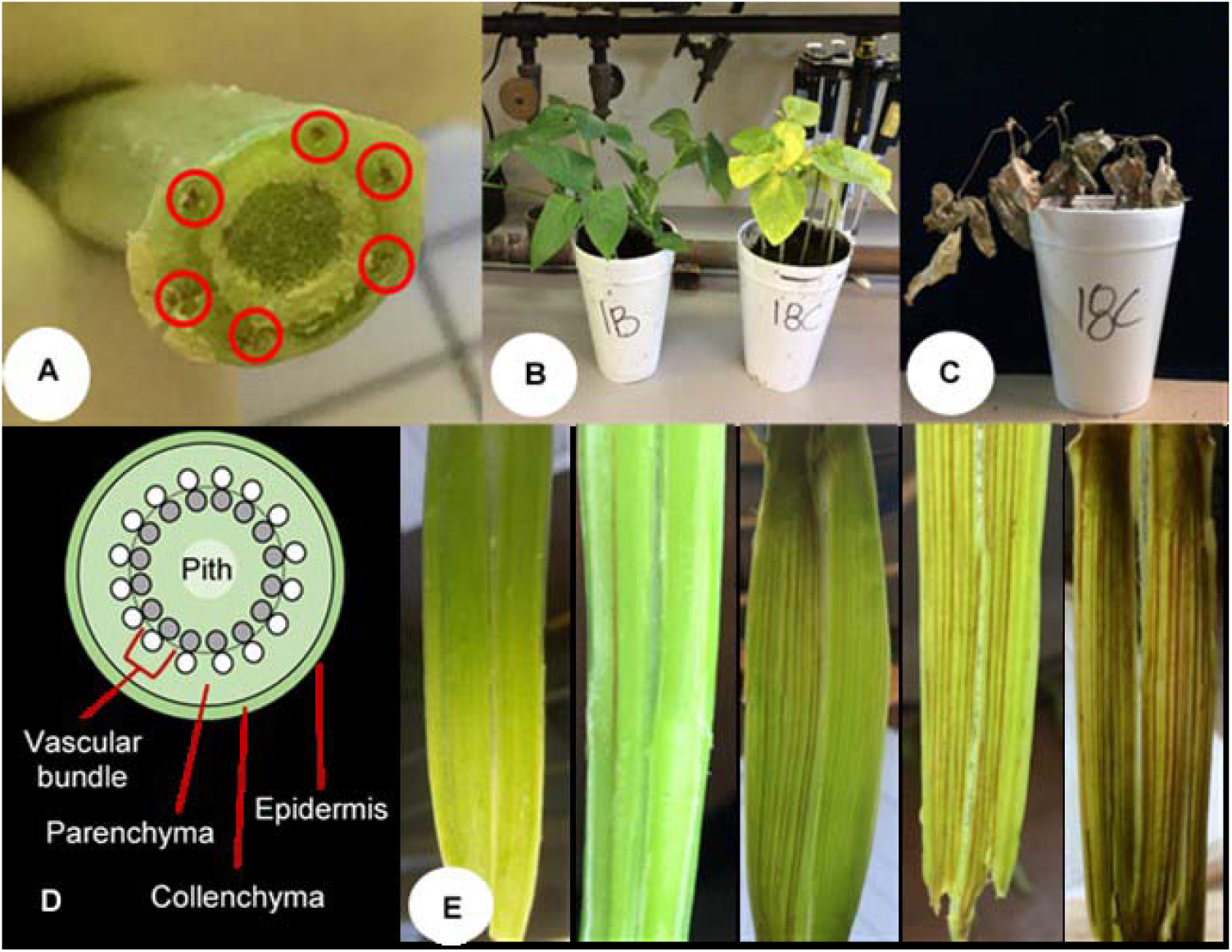
Symptoms of Fusarium wilt incited by *Fusarium oxysporum* f. sp. *tracheiphilum* race 4 (Fot4) inoculated on cowpea plants. **A**, resistant and susceptible plants (pot 1B and 18C, respectively) 20 days after inoculation; **B**, susceptible plants 35 days after inoculation; **C**, cross-section of cowpea plant stem showing vascular discoloration highlighted in circles 35 days after inoculation; **D**, schematic of stem anatomy of a typical dicotyledon plant; gray and white circles = xylem and phloem, respectively; **E**, longitudinal stem sections of cowpea F_2_ lines segregating for resistance 35 days after inoculation; the stems on the far left and far right are of resistant and susceptible plants showing no (left) and severe (right) vascular discoloration, respectively.

### Evaluation of wilting

Thirty-five days after inoculation, individual plant stems were cut at the soil line (Fig. 1A), and the plants evaluated for wilting, %VDL, NFNV, plant height (PH) and shoot-weight (SW). Wilting severity was assessed following a 0 – 5 categorical rating scale (wilt score) modified from Rigert and Foster (1987), where 0 = a healthy plant with no yellowing/wilting symptoms (Fig. 1B – pot 1B); 1 = >0 - 10% of the plant canopy showing yellowing/wilting symptoms (canopy loss up to 10%); 2 = >10 - 25% of the canopy showing yellowing/wilting/stunting (canopy loss more than 10 to 25%); 3 = >25 - 50% yellowing/wilting and canopy loss of more than 25 to 50%, with a dark-brown spot visible on the base of the petiole and on the petiole scar on the stem; 4 = >50 - 75% yellowing/wilting, severe canopy loss (more than 50 to 75%), with the stem colored green-brownish; and 5 = >75 - 100%, the canopy almost completely to completely lost (more than 75 to 100%), the plant is moribund or dead, with limited green-brownish stem to all brown stem tissue, respectively (Fig. 1C).

### Vascular discoloration length (%VDL)

The stem of each plant assessed for wilting was slit open longitudinally with a razor blade from the base to the apex (Fig. 1E), and the extent of vascular discoloration from the base was measured using a ruler. The severity of vascular discoloration length (%VDL) was calculated as the ratio between the length of vascular discoloration and the total plant stem height, 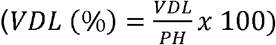.

### Vascular necrosis

After plants were evaluated for wilting and vascular discoloration length, the number of dark-brown discolored vessels (Figs. 1A and 1E) were enumerated in the longitudinally cut stem sections (Fig.1E) to assess the severity of vascular necrosis caused by Fusarium wilt. Enumeration of necrotic vessels on longitudinal cut stem sections is preferable, because on transverse cut stem sections (Figs. 1A) it is hard to visualize clustered-adjacent necrotic vessels. To enumerate necrotic vessels, entire plant stems were cut open along one side, and the stem opened (like opening a book), flattened, placed interior surface up on a porta-trace light box (Gagne Associates, Inc., Binghamton, N. Y.) and the dark-brown vessels counted. In plants with thicker stems, the parenchyma was lightly scalped using a razor blade to facilitate light penetration for better visualization and enumeration of clustered necrotic vessels.

### Data analysis

All analyses were performed using SAS studio version 3.7 (SAS University Edition, SAS, Cary, NC). The association between the plant wilting (wilt score) and vascular discoloration (%VDL), plant wilting and number of Fusarium necrotic vessels (NFNV); and %VDL and NFNV was explored using correlation analysis. Specifically, Pearson’s correlation analysis was performed to explore for the association between %VDL and NFNV since both are continuous variables. The relationships between plant wilting and NFNV and between plant wilting and %VDL were explored using Spearman’s rank correlation because these relationships involved a categorical and a continuous variable.

The relationships between plant wilting (wilt score) and %VDL, and between plant wilting and NFNV were determined using linear regression analysis. Plant wilting (the dependent variable) was regressed against either %VDL or NFNV since plant wilting is a result of vascular occlusion and damage to the plant vascular system (the independent variables were %VDL and NFNV) caused by Fusarium infection. The accuracy of both plant wilting predictors, %VDL and NFNV, were compared using estimated coefficients of determination (*R*^*2*^) from both regression analyses. Although no cause-effect relationship is expected between %VDL and NFNV, regression analysis was performed to examine their relationship in expressing the severity of vascular damage caused by Fusarium wilt and to corroborate the results of the correlation analysis between these phenotypes.

In addition, plant wilting, %VDL and NFNV phenotypic data were tested for several genetic models of segregation between susceptibility – resistance phenotypes to examine segregation distortions using Chi-square goodness-of-fit analysis (Little and Hills, 1978). The Chi-square values were adjusted following Yates correction for continuity (Little and Hills, 1978). The genetic models tested for segregation included a single gene, two dominant genes, and dominant-recessive genes, respectively. The expected segregation ratios between susceptible and resistant phenotypes would be 3:1, 15:1 (9:7) and 13:3, respectively. The ratio 9:7 differs from 15:1 in that it includes the interaction between both genes. These segregation ratios were considered based on reports in the literature describing inheritance of resistance to Fusarium wilt in cowpea and other crops, which is controlled by only a few genes (Netzer et al., 1977; Rigert and Foster, 1987; Salgado et al., 1995; Cross et al., 2000; Pottorff et al., 2014). The distinction between susceptible and resistant phenotypic responses was based on the average plant wilt score, %VDL and NFNV of parents and all populations, and the threshold was set at a plant wilt score of < 3 and ≥ 3 (corresponding values for %VDL and NFNV were used as thresholds for these metrics), which indicate plant wilting of ≤ 25% and plant wilting > 25%, respectively.

## RESULTS

### Association and relationship between wilt phenotype variables

All four susceptible parents (CB46, INIA-73, 24-125B-1 and Bambey-21) developed extensive vascular discoloration, with numerous necrotic vessels, wilted and eventually died following infection with Fot4. In contrast, the resistant parents IT93K-503-1, Ecute and FN-2-9-04 did not exhibit noticeable wilting symptoms (Table 2). However, very few vascular vessels (1 to 3) were detected with necrotic symptoms extending 3 to 26% of the plant height.

**TABLE 2.**
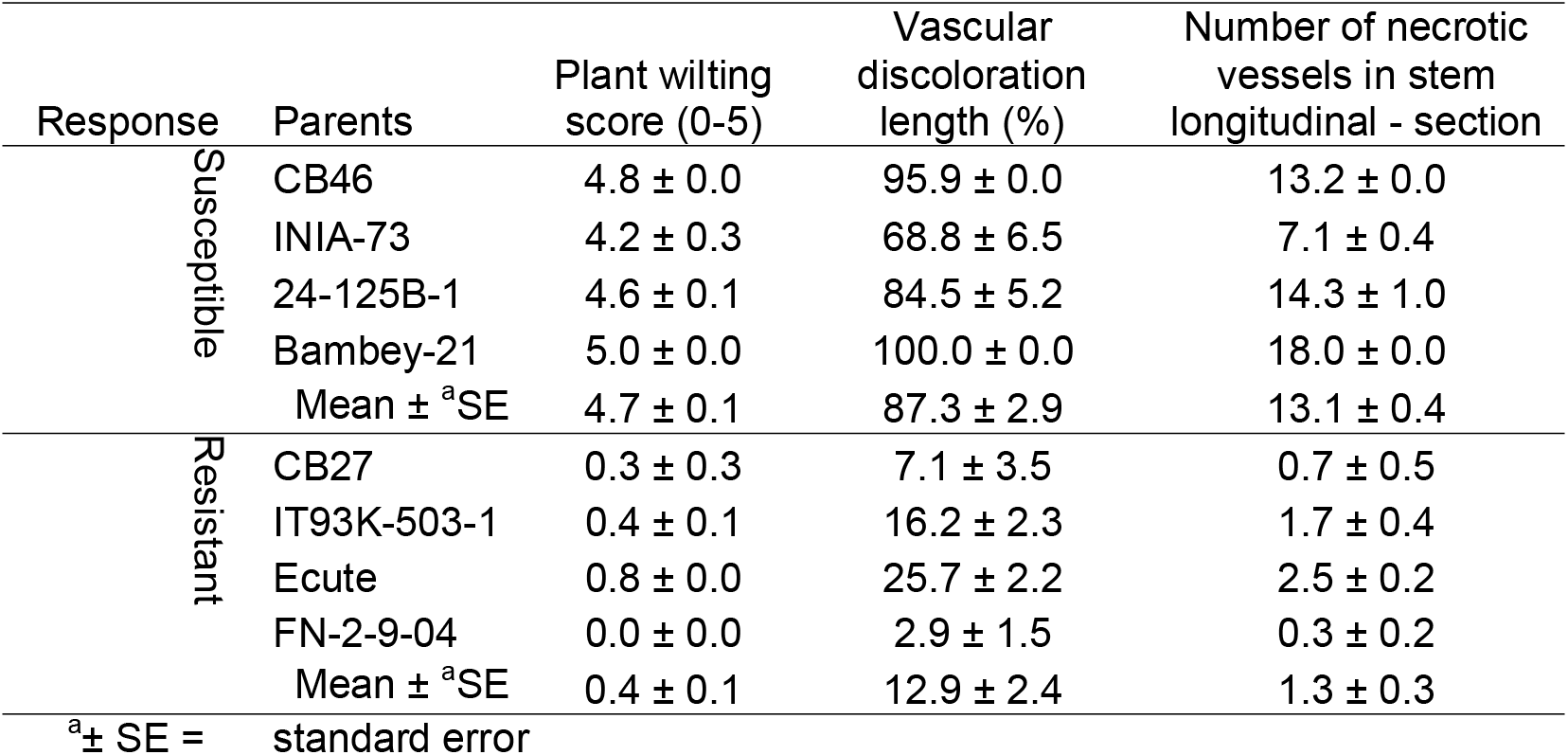
Symptom development in parental genotypes (controls) of cowpea including wilting and development of vascular necrosis induced by *Fusarium oxysporum* f. sp. *tracheiphilum* race 4 (Fot4).

### Correlation analysis

There was strong association among all the variables in the different populations. The correlation coefficients for the association between the severity of plant wilting and NFNV for populations 1, 2, 3, 4, 5, 6 and 7 were 0.874, 0.892, 0.838, 0.941, 0.799, 0.781 and 0.522 (all with *P* < 0.00001), respectively. Correlation coefficients for the association between the severity of plant wilting and %VDL for populations 1, 2, 3, 4, 5, 6 and 7 were 0.892, 0.894, 0.855, 0.922, 0.779, 0.769, and 0.513 (all with *P* < 0.00001), respectively. Correlation coefficients for the association between NFNV and %VDL for populations 1, 2, 3, 4, 5, 6 and 7 were 0.898, 0.913, 0.968, 0.965, 0.940, 0.916 and 0.884 (all with *P* < 0.00001), respectively. Similarly, with the F_2:3_ population comprising 175 families, correlation coefficients of the associations between the severity of plant wilting and NFNV, and between the severity of plant wilting and %VDL; and between NFNV and %VDL were 0.875, 0.925 and 0.832 (all *P* < 0.00001), respectively.

### Regression analysis

In all F_2_ population types (populations 1, 2, 3, 4, 5, 6 and 7), the severity of plant wilting was linearly related to NFNV (Figs. 2A – 2G) and %VDL (Figs. S1A – S1G). For NFNV, the coefficient of determination indicated a moderately strong to strong linear relationship (*R*^*2*^ = 0.78 – 0.91). Similarly, the linear relationship between %VDL and plant wilting was moderately strong to strong (*R^2^ =* 0.81 – 0.87). For all F_2_ populations, there was a consistently strong linear relationship between %VDL and NFNV (*R*^*2*^ = 0.81 – 0.94). Similarly, for the F_2:3_ population comprising 175 families, the linear relationships were strong between the severity of plant wilting and NFNV, and between the severity of plant wilting and %VDL, and between NFNV and %VDL (*R*^*2*^ = 0.71, 0.87 and 0.69, respectively) (Figs. S3A, B and C, respectively).

**Fig. 2.**
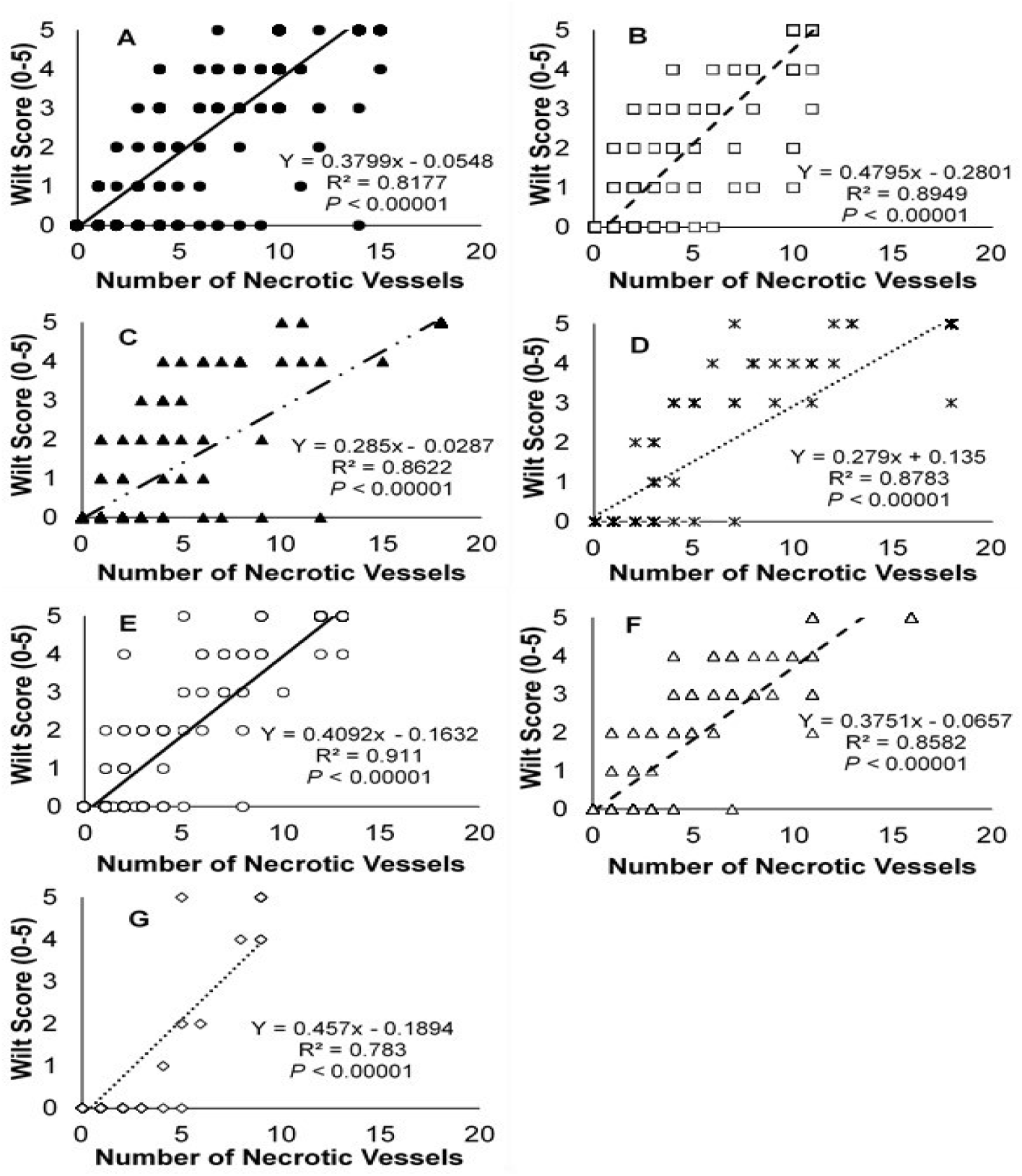
Relationship between wilt score and number of necrotic vessels in seven F_2_ populations (A, B, C, D, E, F and G). **A**, **B**, **C** and **D** (populations 1, 2, 3 and 4) – susceptible × resistant crosses; and **E**, **F** and **G** (populations 5, 6 and 7) – resistant × resistant crosses. The linear regression models describe the dependence of plant wilt on the number of necrotic vessels. The coefficient of determination, *R*^*2*^ describes the proportion of variance of plant wilting explained by the number of necrotic vessels. The *P*-value indicates whether the regression model was significant.

### Plant selection

Thresholds for plant selection between resistant and susceptible plants were determined for NFNV and %VDL using the association of both phenotypes with plant wilt. Plants with NFNV phenotypes ranging from 1 to 18 in the F_2_ populations (1 to 26 for the F_2:3_ population) had corresponding associated severities of wilt. The threshold for resistant and susceptible wilt phenotypes was fixed at disease score < 3 and ≥ 3, respectively. This threshold of wilt symptoms was determined based on the average plant wilting across all populations and parental phenotypes. On the wilt rating scale, a score of 3 indicates plants with > 25 to 50% yellowing/wilting and > 25 to 50% of the canopy wilted or lost. Based on this wilting threshold, critical threshold values were determined for NFNV and %VDL, equivalent to visual estimates of the severity of wilting (Tables 3 and 4).

The F_2_ plants (populations 1, 2, 3, 4, 5, 6 and 7) barely exhibiting visible wilt symptoms the mean wilt score was 0.1 to 0.5, or less than 10% of the plant wilted. The NFNV score for these plants ranged from 1 to 3, and the mean %VDL from 13.2 to 25.2%. Plants with a mean wilt score of 1.6 - 2.1 (approximately > 10 to 25% of the plant wilted) had an NFNV of 4 to 5, which corresponded to a mean %VDL of 29.6 – 32.7% (Table 3). Wilting of 50% (wilt score of 2.8 - 2.9) was observed in plants with an NFNV of 6 or 7, and a corresponding mean %VDL of 45.2 to 49.0%; whereas plants ≥ 75% wilted (a wilt score of 3.4 to 4.6) had an NFNV ≥ 8, which corresponded to a %VDL of 46.1 to 76.7%. The total number of vessels in each plant was not enumerated, but it was observed that even the highly susceptible genotypes with 10 to 18 necrotic vessels in the F2 generation, had some vessels that were not necrotic (Table 3).

**TABLE 3.** The number of Fusarium necrotic vessels (NFNV), the plant wilt symptom score and the extent of vascular discoloration (%VDL) observed among seven F_2_ populations of cowpea inoculated with *Fusarium oxysporum f. sp. tracheiphilum* race 4 (Fot4).

| Variable | <sup>b</sup>Pop. | NFNV |  |  |  |  |  |  |  |  |  |
| --- | --- | --- | --- | --- | --- | --- | --- | --- | --- | --- | --- |
|  |  | 1 | 2 | 3 | 4 | 5 | 6 | 7 | 8 | 9 | 10 - 18 |
|  |  | <sup>a</sup>Number of plants |  |  |  |  |  |  |  |  |  |
|  |  | 31-68 | 7-42 | 4-26 | 2-23 | 3-11 | 1-8 | 2-12 | 1-10 | 2-6 | 42-128 |
| Plant wilting<br>(score, 0-5) | 1 | 0.2 | 0.2 | 0.6 | 1.9 | 0.9 | 2.3 | 2.9 | 2.9 | 2.8 | 4.5 |
|  | 2 | 0.1 | 0.5 | 0.7 | 1.8 | 1.6 | 2.6 | 2.8 | 3.0 | <sup>c</sup>- | 4.7 |
|  | 3 | 0.2 | 0.2 | 0.4 | 1.7 | 2.5 | 2.2 | 2.0 | 4.0 | 1.0 | 4.5 |
|  | 4 | 0.0 | 0.3 | 0.6 | 2.0 | 2.3 | 4.0 | 2.8 | 4.0 | 3.5 | 4.0 |
|  | 5 | 0.1 | 0.3 | 0.6 | 1.1 | 2.5 | 3.3 | 3.5 | 2.6 | 4.5 | 4.9 |
|  | 6 | 0.1 | 0.2 | 0.5 | 2.0 | 2.5 | 3.3 | 3.1 | 3.3 | 3.5 | 4.7 |
|  | 7 | 0.0 | 0.0 | 0.0 | 0.5 | 2.3 | 2.0 | <sup>c</sup>- | 4.0 | 4.7 | <sup>c</sup>- |
|  | Mean | 0.1 | 0.3 | 0.5 | 1.6 | 2.1 | 2.8 | 2.9 | 3.4 | 3.3 | 4.6 |
|  | Standard Error | 0.0 | 0.1 | 0.1 | 0.2 | 0.2 | 0.3 | 0.4 | 0.2 | 0.5 | 0.1 |
| %VDL | 1 | 16.1 | 18.7 | 27.8 | 34.4 | 32.3 | 44.6 | 51.1 | 43.8 | 44.3 | 88.5 |
|  | 2 | 18.9 | 25.9 | 28.3 | 34.7 | 41.5 | 45.0 | 45.6 | 38.8 | <sup>c</sup>- | 91.5 |
|  | 3 | 11.4 | 14.0 | 24.8 | 24.6 | 29.2 | 31.2 | 46.0 | 42.5 | 32.4 | 58.4 |
|  | 4 | 7.3 | 16.6 | 21.6 | 28.2 | 33.9 | 69.5 | 46.8 | 58.9 | 42.7 | 34.8 |
|  | 5 | 19.5 | 26.5 | 30.0 | 38.8 | 2.5 | 56.9 | 49.9 | 61.2 | 75.4 | 97.6 |
|  | 6 | 14.5 | 19.4 | 29.2 | 34.5 | 41.3 | 48.2 | 54.3 | 45.6 | 43.7 | 89.3 |
|  | 7 | 4.7 | 8.9 | 14.7 | 11.8 | 47.9 | 20.8 | <sup>c</sup>- | 32.0 | 79.0 | <sup>c</sup>- |
|  | Mean | 13.2 | 18.6 | 25.2 | 29.6 | 32.7 | 45.2 | 49.0 | 46.1 | 52.9 | 76.7 |
|  | Standard Error | 2.1 | 2.4 | 2.1 | 3.5 | 5.6 | 6.0 | 7.1 | 4.0 | 7.3 | 13.9 |

<sup>a</sup>The number of plants identified within each NFNV varied among seven populations, and each NFNV was associated with a wilt score and extent of vascular discoloration in each population. The NFNV among populations varied from 1 to 18.
<sup>b</sup>Pop = populations, indicate in the second column; Pop 1 = CB46/FN-2-9-04, 2 = INIA-73/FN-2-9-04, 3 = 24-125B-1/FN-2-9-04, 4 = Bambey-21/FN-2-9-04, 5 = CB27/FN-2-9-04, 6 = IT93K-503-1/FN-2-9-04 and 7 = Ecute/FN-2-9-04.
<sup>c</sup>indicates no plant was identified within that NFNV class.

Also, the F_2:3_ population (population 8), which comprised 175 families (on average 25 plants/family) showed evidence of associations among assessment methods. Plants with mean wilt scores of 0.3 to 1.1 (> 0 to 10% wilted) had an NFNV of 1 – 3 and a mean %VDL ranging from 5.5 to 16.7% (Table 4). Plants with wilt scores of 1.6 to 2.1 (> 10 to 25% wilted) had an NFNV of 4 to 6 and a mean %VDL ranging from 27.1 to 36.2, respectively. Plants that were 50% wilted (wilt score of 2.5 to 3.2) had an NFNV of 7 to 9, which corresponded to a mean %VDL of 46.1 to 55.4%, respectively. Plants with wilt score of 3.8 (≥ 75% wilted) had and NFNV ≥ 10, and a %VDL ≥ 69.9.

**TABLE 4.**
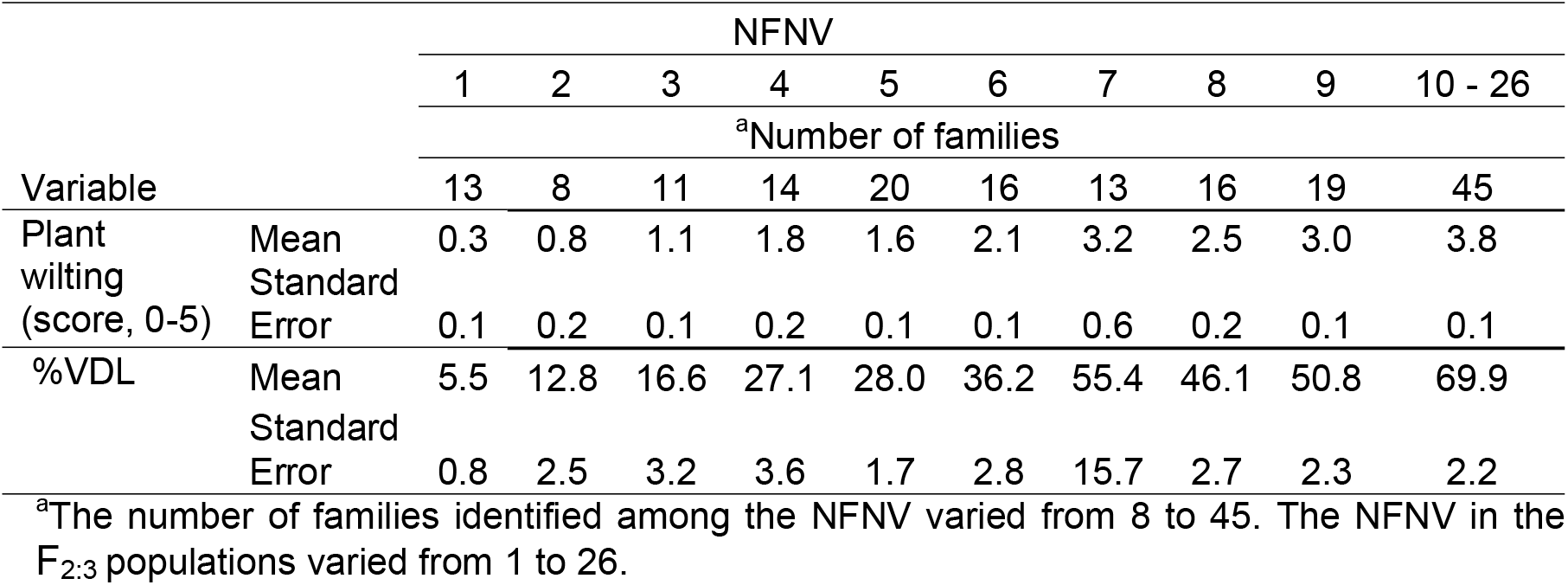
The number of Fusarium necrotic vessels (NFNV), the plant wilt symptom score and the extent of vascular discoloration (%VDL) observed in the F_2:3_ population (Population 8, Table 1) of cowpea inoculated with *Fusarium oxysporum f. sp. tracheiphilum* race 4 (Fot4).

### Segregation distortion

Two F_2_ populations (susceptible × resistant population 1 and resistant × resistant population 5) were analyzed to determine the levels of segregation distortion observed for each Fusarium wilt severity assessment metric: wilting, vascular discoloration length (%VDL) and number of Fusarium necrotic vessels (NFNV). A cut-off between resistant and susceptible phenotypes was set at a wilt score of < 3, a %VDL < 45 and an NFNV count < 6. These thresholds were based on 50% plant wilt and the phenotypic responses of the F_2_ population to wilting, vascular discoloration length and vascular necrosis (Table 3).

The best fitting segregation ratio (Resistant:Susceptible) for wilting (206:158), %VDL (225:139) and NFNV (208:156) in F_2_ population 1 was 9:7 (Table 5); however, this was significant only for wilt score (*X^2^ =* 0.00*, P =* 0.99) and NFNV (*X^2^ =* 0.08*, P =* 0.75 – 0.90). Segregation distortion was higher for %VDL compared to the wilt score and NFNV, and the differences between observed and expected phenotypic responses were 19.75, 0.75 and 2.75, respectively. In the resistant × resistant F_2_ population 5, the best fit segregation ratio for wilt score (251:72), %VDL (242:81) and NFNV (251:72) was 13:3. This model was significant for the wilt score and NFNV (*X*^*2*^ = 2.43, *P* = 0.10 – 0.25 for both phenotypic metrics), but it was not significant for %VDL (*X*^*2*^ = 8.08, *P* < 0.01). Also, in F_2_ population 1, both wilt score and NFNV had lower segregation distortion (10.94) compared to %VDL (19.94).

**TABLE 5.** Segregation for wilting (wilt score), vascular discoloration length (%VDL) and number of Fusarium necrotic vessels (NFNV) in two F_2_ populations (population 1 = susceptible × resistant; population 5 = resistant × resistant) of cowpea inoculated with *Fusarium oxysporum f. sp. tracheiphilum* race 4 (Fot4)

|  |  | Population 1 |  |  |  | Population 5 |  |  |  |
| --- | --- | --- | --- | --- | --- | --- | --- | --- | --- |
| Variable | Response | <sup>a</sup>O | <sup>b</sup>E (9:7) | <sup>c</sup>O-E | <sup>d</sup>X<sup>2</sup> | <sup>a</sup>O | <sup>b</sup>E (13:3) | <sup>c</sup>O-E | <sup>d</sup>X<sup>2</sup> |
| Wilt score | Resistant | 206 | 204.75 | 0.75 | 0.00 | 251 | 262.44 | 10.94 | 0.46 |
|  | Susceptible | 158 | 159.25 | 0.75 | 0.00 | 72 | 60.56 | 10.94 | 1.98 |
|  | Total | 364 | 364 | 1.5 | 0.00 | 323 | 323 | 21.88 | 2.43 |
| %VDL | Resistant | 225 | 204.75 | 19.75 | 1.91 | 242 | 262.44 | 19.94 | 1.51 |
|  | Susceptible | 139 | 159.25 | 19.75 | 2.45 | 81 | 60.56 | 19.94 | 6.56 |
|  | Total | 364 | 364 | 39.50 | 4.35 | 323 | 323 | 39.88 | 8.08 |
| NFNV | Resistant | 208 | 204.75 | 2.75 | 0.04 | 251 | 262.44 | 10.94 | 0.46 |
|  | Susceptible | 156 | 159.25 | 2.75 | 0.05 | 72 | 60.56 | 10.94 | 1.98 |
|  | Total | 364 | 364 | 5.50 | 0.08 | 323 | 323 | 21.88 | 2.43 |

<sup>a</sup>O and <sup>b</sup>E = observed and expected number of resistant or susceptible plants (values in brackets are the significant expected model ratios between resistant and susceptible plants);
<sup>c</sup>O-E = deviation from expected value;
<sup>d</sup>X<sup>2</sup> = chi-square; E (R:S) = expected model ratio.

## DISCUSSION

The plant wilt score was highly correlated with both vascular discoloration length (%VDL) and number of necrotic vessels (NFNV) in the seven F_2_ populations and one F_2:3_ population of cowpea. Associations between wilting and vascular discoloration induced by Fusarium has been reported in tomato, common bean and cotton (Gao et al., 1994; Buruchara and Camacho, 2000; Ulloa et al. 2006); however, in soybean, these phenotypic responses were not associated with one another (Luckew et al., 2012). In this study, wilt response was correlated with both %VDL and NFNV, and most of the variability in plant wilt phenotypes was accounted for by both %VDL and NFNV, as indicated by the coefficients of determination (*R*^*2*^ all ≥ 0.71); however, the use of only one of the three phenotypes (plant wilt, %VDL or NFNV) alone may not be completely effective in measuring disease severity in the cowpea – *Fusarium* pathosystem and quite possibly in other plant wilt pathosystems. Other factors including germplasm and pathogen isolate origin, the anatomy of the vascular system and measurement errors may undermine the relationship, as indicated by segregation distortion. For example, in studies on Fusarium wilt of cotton, phenotyping based on a combination of plant wilting and %VDL has proved effective (Ulloa et al., 2006; Wang et al., 2017). In cowpea, recently we have demonstrated through a series of genetic studies and quantitative trait loci mapping that determinants for resistance to wilt caused by *Fusarium oxysporum* f. sp*. tracheiphilum* race 4 in accession FN-2-9-04 (the resistant parent used to develop populations used in this study) are located on two chromosomes, and resistance to %VDL and NFNV are co-located with resistance to wilting on only one of the chromosomes (Ndeve, 2017). This finding indicated that plant wilting may be a complex trait under control by several genes, some of which may be associated with resistance to vascular discoloration and to the proliferation in the number of necrotic vessels, which are associated with the development of the wilting phenotype.

The phenotypes %VDL and NFNV were highly correlated; however, %VDL expresses the extent of vertical vascular discoloration in the plant stem, but it does not take into account the number of necrotic vessels in the plant stem required to trigger the plant wilt symptoms resulting from occluded and damaged vessels. For example, in some cases, infected plants have one to four discolored vessels extending >50% of the plant height, but they will have no visual symptoms of wilt; therefore, %VDL will not be correlated with plant wilt phenotype and %VDL measurements will overestimate the severity of vascular damage. In this situation, it is important to determine the number of discolored vessels for accurate phenotype-based plant selection. Vascular discoloration length (%VDL) and NFNV were contrasted to determine how closely their values reflect the severity of plant wilt (True value) and their usefulness for plant selection for resistance to Fusarium wilt disease. The analyses showed that in both the F_2_ and F_2:3_ test populations a vascular discoloration length of 45% was associated with a wilt score of 3 (50%), and this wilt score was associated with a count of 6 to 7 necrotic vessels. Both the %VDL and NFNV are quantitative metrics that provide objective measurement (as opposed to a subjective estimate) of Fusarium wilt severity in cowpea plants, and the data can be analyzed using parametric statistics (Campbell and Neher, 1994). Conversely, rating the extent of vascular discoloration and wilting symptoms using a categorical scale (e.g., 0-5) with corresponding disease severity classes indicating percent vascular discoloration or percent of plant wilting (0, 1, 2, 3, 4 and 5 = 0, 10, 25, 50, 75 and 100%) (Pottorff et al., 2012; Pottorff et al., 2014) can lead to errors in disease assessment due to subjectivity inherent among raters, which can be exacerbated by inexperienced or less accurate disease evaluators (Nutter et al., 1993; Bock et al., 2010). Visual assessment of the severity of Fusarium wilt symptoms relies on visual rating scales that express the severity of plant wilting, and the externally visible symptoms of wilt are a direct consequence of damage to the vascular system. The modifications to the process of Fusarium wilt disease assessment described in this study allow for quantitative measurement of severity of vascular colonization by a combination of length of vascular discoloration (%VDL) and enumeration of necrotic vessels (NFNV). Using NFNV was less error prone compared to measuring the length of vascular discoloration, as indicated by the segregation distortion between the observed and expected numbers of resistant and susceptible plants. Because symptoms of Fusarium wilt-induced vascular damage are generally similar in a range of crop plant species, including cotton (Ulloa et al., 2006; Wang et al., 2017) and tomato (Buruchara and Camacho, 2000), the protocol described in this study may be usefully applied to assess Fusarium wilt disease severity on these crops and in other plant-Fusarium wilt pathosystems. Enumerating the number of necrotic vessels in infected plants is time consuming, which is an important consideration in breeding programs where phenotyping large numbers of plants is required. But both NFNV and %VDL may be amenable to measurement using digital-image phenotyping technology (Nutter and Schultz, 1995; Bock and Nutter, 2011; Barbedo, 2013; Mutka and Bart, 2015; Mahlein, 2016). Image analysis software can be programmed to count the number and the length of vascular necrotic vessels (NFNV and %VDL) in images of longitudinal plant sections. Although this method has not been explored it could reduce the labor required and may provide added accuracy and repeatability of disease severity estimates in Fusarium wilt phenotyping, which are crucial for effective plant selection and breeding for Fusarium wilt resistance.

The accuracy and precision of digital-image based plant disease phenotyping varies with pathosystem (Bock and Nutter, 2011; Mutka and Bart, 2015). For example, measurements of disease severity of sunflower blight and oat leaf rust using digital-image analysis were overestimated compared to visual estimates (Tucker and Chakraborty, 1997). Similar results were reported by Olmstead et al. (2001) when assessing powdery mildew disease severity on sweet cherry, whereas radiometric assessment of dollar spot severity on bentgrass was more precise than visual assessment (Nutter et al., 1993). Bock et al. (2008) reported that digital-image phenotyping of citrus canker on grapefruit leaves was more consistent and accurate when compared to visual disease assessment. Thus, the accuracy, precision, repeatability and reproducibility of image-based plant disease assessments varies among pathosystems and are likely affected by the symptoms, characteristics of the image, and the conditions under which the images were captured (Bock and Nutter, 2011).

This study was based on F_2_ and F_2:3_ populations because the segregation and zygosity at these generations enabled a broad assessment of the variability of the phenotypic variables of interest, which are less apparent in advanced highly inbred lines (e.g. F_8_ or F_10_). In plant breeding programs, plant selections for wilt resistance traits must be strategized to avoid loss of breeding material to Fusarium wilt infection, including through destructive assessments, such as the ones described in this study. In early generations, for example F_2_ populations, plants are not inoculated with Fusarium wilt and leaf samples can be collected for DNA extraction and subsequent genotyping using previously characterized molecular markers (Ndeve, 2017). Each F_2_ plant is managed to produce enough seed for the next generation (F_2:3_) for conducting further Fusarium wilt resistance phenotyping. After phenotyping the F_2:3_ for plant wilting, %VDL and/or NFNV, the phenotypic data are associated with genotypic data through marker-trait association analysis. This approach associates the plant phenotype (plant wilting, %VDL and/or NFNV) with the specific target trait alleles to allow the identification and selection of plants of interest (these materials trace back to their F_2:3_ seed stocks). Also, this approach enables location of the resistance gene(s) associated with the phenotypes in the plant genome and reduces the time required for the breeding cycle. Since all phenotyped F_2:3_ plants are either killed by Fusarium wilt disease or destroyed during plant phenotyping, the data from marker-trait association analysis enables tracing the identity and selection of plants of interest (resistant) back to their original seed stocks, which can be used for subsequent advancement of selected breeding material. Similar approaches using phenotyping and genotyping on different generations (Zhang and Xu, 2004) can be employed including marker-assisted backcrossing and marker-assisted recurrent selection. In this study, the variation in phenotypic response (wilt score, NFNV and %VDL) was a desirable condition in that it allowed detection of extreme phenotypes (parental), and intermediate phenotypes resulting from quantitative inheritance and heterozygosity. In segregating breeding populations this phenotypic variation provides a basis for understanding the genetic control of the traits including the number, dominance and additive effects of the genes involved.

This study introduces the number of Fusarium necrotic vessels (NFNV) as a novel quantitative metric to measure the severity of vascular damage caused by Fusarium wilt disease of cowpea; the strong correlation of NFNV with vascular discoloration length (%VDL), a metric used to measure the severity of vascular damage, showed that both metrics provide accurate measurements of the severity of plant vascular damage under Fusarium wilt disease infection. Also, the strong correlation between NFVN and %VDL indicated that either of these metrics can be used to phenotype plant vascular damage; however, when using %VDL to phenotype vascular damage, plants showing one to four discolored vessels that extend > 50% of the plant height may cause segregation distortion of the ratio between the observed and expected number of plants resistant and susceptible to vascular damage, respectively. In addition, the linear relationship between both vascular metrics and plant wilting phenotype, also provided evidence of the accuracy of NFVN and %VDL as a gauge of the severity of vascular damage and to predict plant wilt phenotype resulting from Fusarium wilt infection. When searching for novel sources of resistance to Fusarium wilt in germplasm collections, it is appropriate to phenotype plants by both plant wilting and vascular damage (NFVN or %VDL) because high phenotypic variation is often present in these materials, allowing identification of resistant, susceptible and partially resistant genotypes. These variations in Fusarium wilt disease phenotypic responses may indicate to some extent a weak correlation between plant wilting and vascular phenotypes. Also, unless the relationship between plant wilting and vascular phenotypes is known in the parental genotypes, plants should be phenotyped for both plant wilt phenotype and vascular phenotype (NFNV or %VDL). Based on this study, both NFNV and %VDL can be used in breeding programs in early or late generation plant phenotyping, providing objective quantification of vascular wilting.

## ACKNOWLEDGEMENTS

Support for this work was provided by grants from the Generation Challenge Program of the Consultative Group on International Agricultural Research, the USAID Dry Grain Pulses CRSP and Innovation Lab for Collaborative Research on Grain Legumes, the California Dry Bean Advisory Board, and the California Agricultural Experiment Station.

---

**S1.**
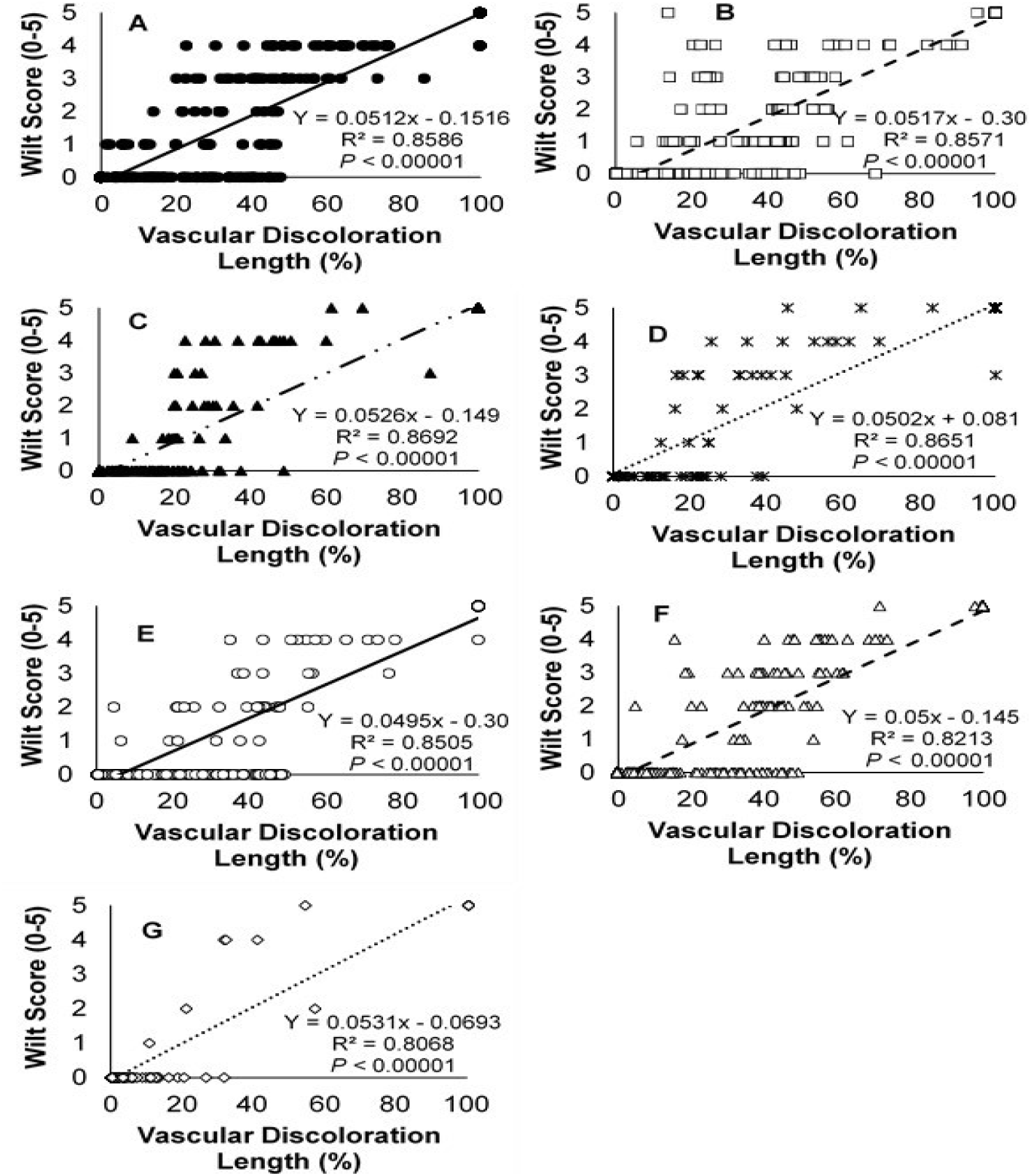
Relationship between wilt score and vascular discoloration length in seven F_2_ populations. **A**, **B**, **C** and **D** (populations 1, 2, 3 and 4) – susceptible × resistant crosses; **E**, **F** and **G** (populations 5, 6 and 7) – resistant × resistant crosses. The linear regression models describe the dependence of plant wilt on vascular discoloration length. The coefficient of determination, *R*^*2*^ describes the proportion of variance of plant wilting explained by the vascular discoloration length. The *P*-value indicates whether the regression model was significant.

**S2.**
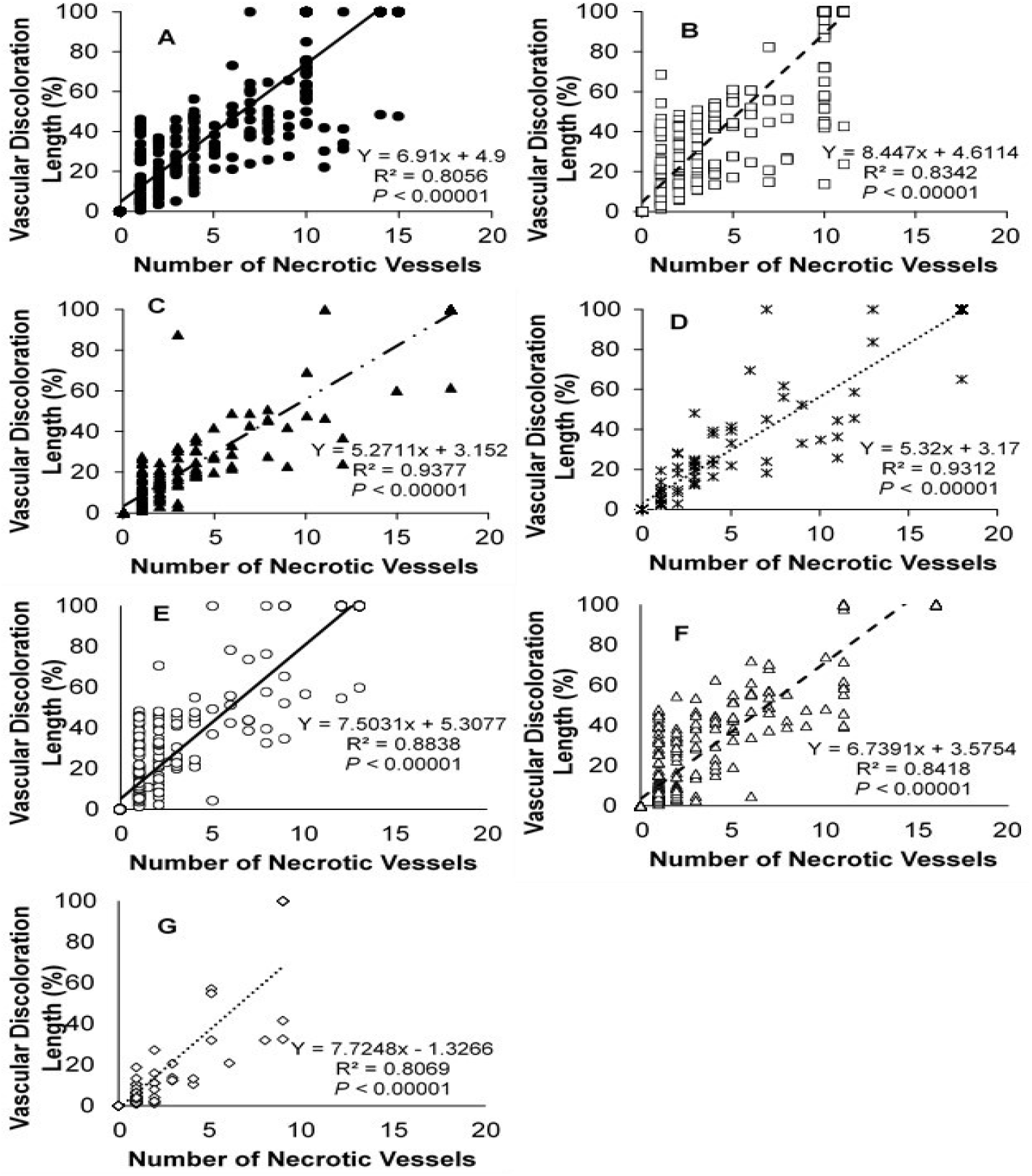
Relationship between vascular discoloration length and number of necrotic vessels in seven F_2_ populations (A, B, C, D, E, F and G). **A**, **B**, **C** and **D** (populations 1, 2, 3 and 4) – susceptible x resistant crosses; **E**, **F** and **G** (populations 5, 6 and 7) – resistant × resistant crosses. The linear regression models describe the relationship between vascular discoloration length and the number of necrotic vessels. The coefficient of determination, *R*^*2*^ describes the proportion of variance of necrotic vessels explained by the vascular discoloration length. The *P*-value indicates whether the regression model was significant.

**S3.**
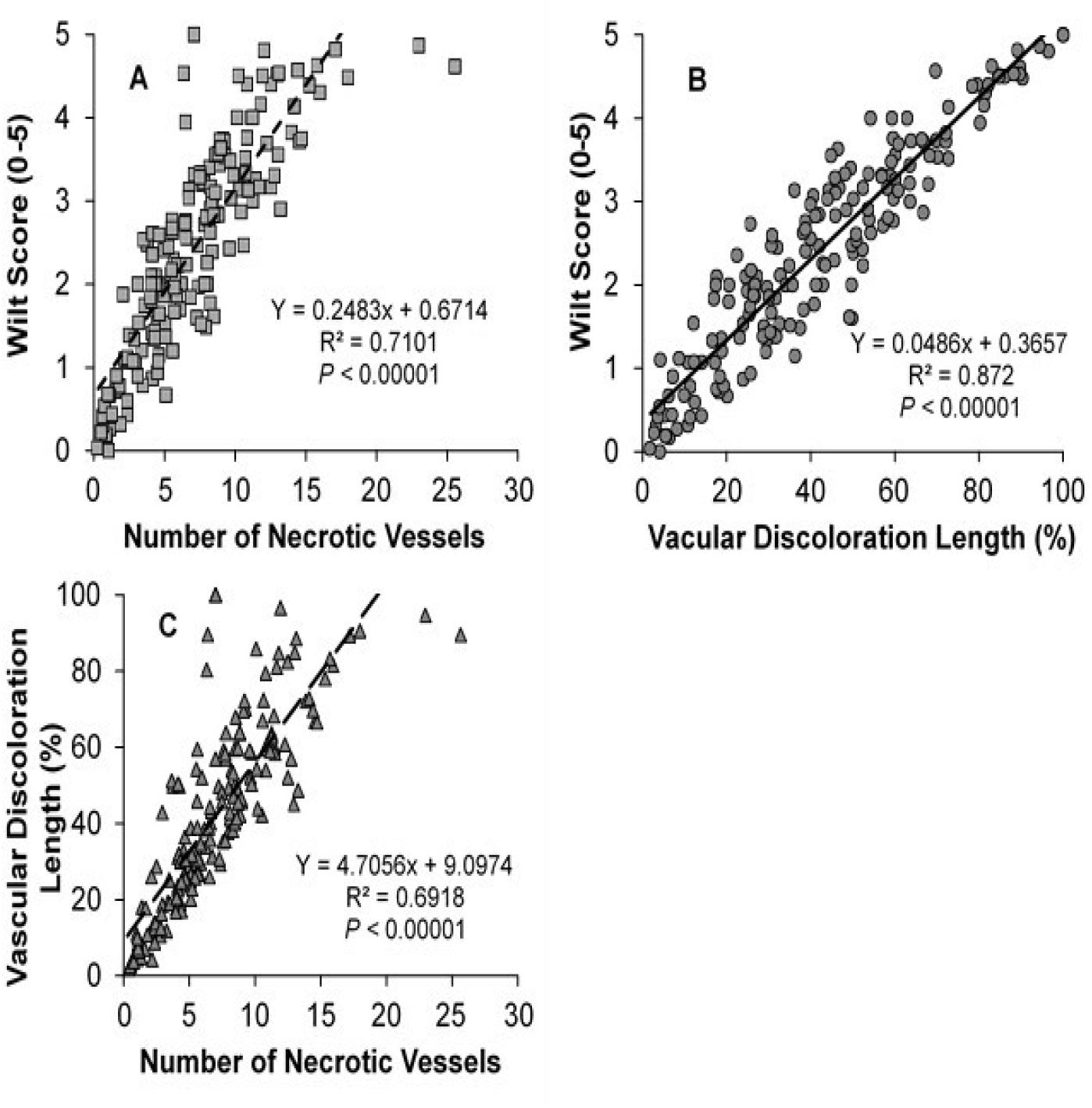
**A** and **B:** The relationship between wilt score and the number of necrotic vessels and vascular discoloration, respectively in the F_2:3_ population (population 8). **C:** the relationship between vascular discoloration and the number of necrotic vessels in the F_2:3_ population. The linear regression models are represented by the curve and equation. The coefficient of determination, *R*^*2*^ explains the proportion of variance of plant wilting explained by **A:** number of necrotic vessels and **B:** vascular discoloration length; and **C:** proportion of variance explained by the relationship between both metrics measures to describe vascular phenotypes. The *P*-value indicates whether the regression model was significant.

